# ADHD genetic liability and physical health outcomes - A two-sample Mendelian randomization study

**DOI:** 10.1101/630467

**Authors:** Beate Leppert, Lucy Riglin, Christina Dardani, Ajay Thapar, James R Staley, Kate Tilling, George Davey Smith, Anita Thapar, Evie Stergiakouli

**Author notes:** **Corresponding author:** Dr. Beate Leppert, MRC Integrated Epidemiology Unit, University of Bristol, Oakfield House, Oakfield Grove, Bristol BS8 2BN, UK.

## Abstract

**Objective:** Attention-deficit/hyperactivity disorder (ADHD) has been associated with a broad range of physical health problems, including cardiometabolic, neurological and immunological conditions. Determining whether ADHD plays a causal role in these associations is of great importance for treatment and prevention but also because comorbid health problems further increase the serious social and economic impacts of ADHD on individuals and their families.

**Methods:** We used a two-sample Mendelian randomization (MR) approach to examine the causal relationships between genetic liability for ADHD and previously implicated physical health conditions. 11 genetic variants associated with ADHD were obtained from the latest summary statistics. Consistent effects obtained from IVW, weighted median and MR Egger methods were taken forward for sensitivity analysis, including bidirectional MR and multivariable MR (MVMR).

**Results:** We found evidence of a causal effect of genetic liability for ADHD on childhood obesity (OR:1.29 (95% CI:1.02,1.63)) and coronary artery disease (CAD) (OR:1.11 (95% CI:1.03,1.19)) with consistent results across different MR approaches. There was further evidence for a bidirectional relationship between genetic liability for ADHD and childhood obesity. The effect of genetic liability for ADHD on CAD was independent of smoking heaviness but was attenuated when simultaneously controlling for childhood obesity. There was little evidence for a causal effect on other cardiometabolic, immunological, neurological disorders and lung cancer.

**Conclusion:** Our findings strengthen the argument for early treatment and support for children with ADHD and their families and especially promoting physical activity and providing them with dietary advice to reduce future risk for developing CAD.

**Key Message:**

- Epidemiological studies have reported observational associations between ADHD and adult onset physical health outcomes.
- Mendelian Randomization can be used to assess causal associations for ADHD on health outcomes that would traditionally require long term follow-up and may suffer confounding
- We found that genetic liability for ADHD was associated with coronary artery disease and there was evidence for a bidirectional association between genetic liability for ADHD and childhood obesity
- Multivariable mendelian randomization suggests that the link between genetic liability and coronary artery disease might partially act through childhood obesity but was independent of smoking heaviness
- There was little evidence of a causal of ADHD on other cardiometabolic and immunological diseases.

## INTRODUCTION

Attention-deficit/hyperactivity disorder (ADHD) is the most common neurodevelopmental condition with an onset typically in early childhood and a worldwide prevalence of about 5% in school-aged children and about 2-3% in adults (1, 2). In 65% of children diagnosed with ADHD symptoms impairment persist into adulthood and can be associated with psychiatric problems, social and occupational difficulties (3). ADHD is associated with higher mortality and morbidity rates(4) and a broad range of physical health problems. These include metabolic problems (including obesity and type 2 diabetes mellitus), cardiovascular problems (such as hypertension(5) and other risk factors for cardiovascular disease (CVD)), neurological problems (including epilepsy and migraine(6)), allergic/inflammatory disorders (including asthma (6–8), allergic rhinitis and coeliac disease)(6) and some types of cancer.

Determining whether ADHD causally contributes to these health problems is of great importance because comorbid health problems further increase the serious social and economic impacts of ADHD on individuals and society (9, 10). However, there are multiple possible explanations for the association between ADHD and physical health problems:

1. ADHD could cause physical health problems by increasing risk for an unhealthy lifestyle and risk behaviours, e.g. via smoking, excess alcohol consumption, sedentary lifestyle and dietary choices all of which appear to influence physical ill-health.
2. ADHD could be associated with physical health outcomes because of shared genetic variants (pleiotropic effects).
3. Shared environmental risk (pre- and postnatal factors) may confound the association between ADHD and poor physical health.

Causal relationships between ADHD and physical health problems cannot be resolved by conventional observational studies because of issues including selection bias, reverse causation and residual confounding (11). However recent advances in genome-wide association studies enable the use of genetic variants strongly associated with ADHD (12) as proxies (instrumental variables) for the life-time risk of ADHD and to study causal effects of genetic liability for ADHD on physical health outcomes using Mendelian randomization (MR). The rationale behind MR is that genetic variants can be used as theoretically un-confounded proxies for that exposure since they are determined randomly at conception and segregated through to viable offspring independently of environmental influences(13, 14).

In this study, we aim to investigate causal relationships between genetic liability for ADHD and metabolic, cardiovascular, inflammatory and neurological conditions previously observed to be associated with ADHD, using a two-sample MR design.

## METHODS

### Two-sample MR

MR utilizes genetic variants as proxies for an exposure of interest and allows the estimation of an un-confounded causal association between exposure and outcome, if certain assumptions(13) hold true:

1. The genetic variants are strongly associated with the exposure of interest.
2. The genetic variants are independent of confounders of exposure and outcome.
3. The genetic variants do not affect the outcome except through the exposure (exclusion restriction criterium). If they affect the outcome through other pathways, this is called horizontal pleiotropy.

Within a two-sample MR approach GWAS summary statistics are used to assess both genetic variant-exposure and genetic variant-outcome associations. All steps of the two-sample MR performed here are explained in more detail below and summarized in Figure 1. Analyses were conducted using the TwoSampleMR package version 0.4.14 for R (version 3.4.1).

**Figure 1.**
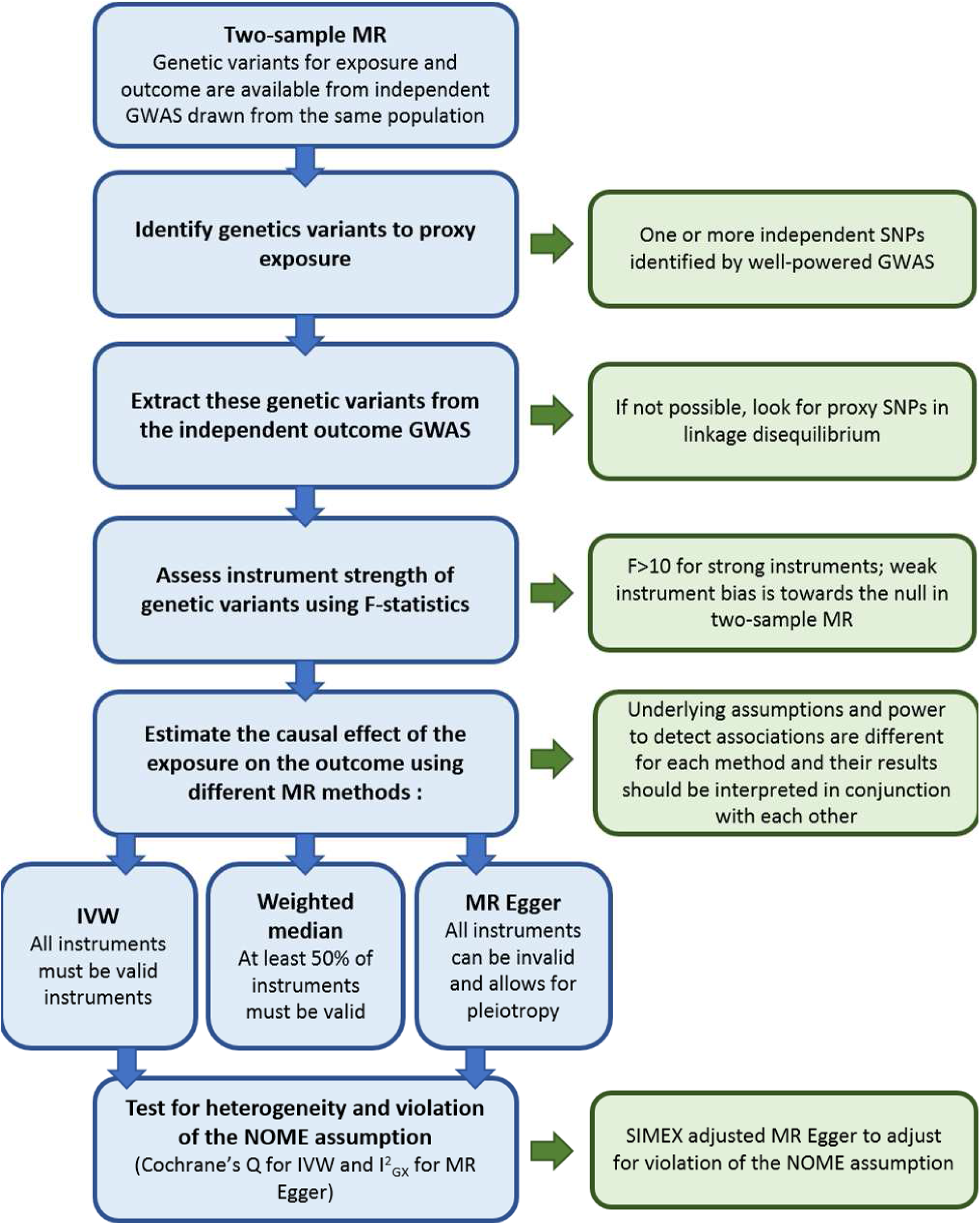
Flow chart of MR study design.

### ADHD GWAS (exposure)

Genetic variants associated with ADHD (the exposure) we identified in the summary statistics for European ancestry participants from the combined PGC+iPSYCH metaanalysis of ADHD including 19 099 cases and 34 194 controls (12). Log odds ratio (OR) and standard error estimates for thirteen SNPs with p<1×10^−7^ were extracted after clumping (r^2^=0.001 within 10,000kb) and removal of palindromic sequences (Table S1 in the Supplement).

### GWAS on physical health problems (outcome)

We used freely available GWAS summary statistics from predominantly European ancestry populations on physical health problems that have been previously reported as being associated with ADHD (6–8). The outcomes included cardiometabolic(15–19), neurological(20), immunological diseases(18, 21–24) and lung cancer (25) (detailed overview in Table 1 and Table S2), because of the strong link between ADHD and smoking behaviour. An exploratory analysis was conducted for metabolic markers derived from the 2016 GWAS from Kettunen et al., since they have been implicated as risk factors for CAD and obesity (26). GWAS derived from UK biobank by the Neale lab (http://www.nealelab.is/uk-biobank) are reported on a risk difference scale and were transformed onto the log OR scale (see supplementary material). All outcome GWAS are thought to be independent of the PGC+iPSYCH sample. The GWAS for coronary artery disease and myocardial infarction are derived from a mixed population sample with 77% white European participants.

**Table 1:**
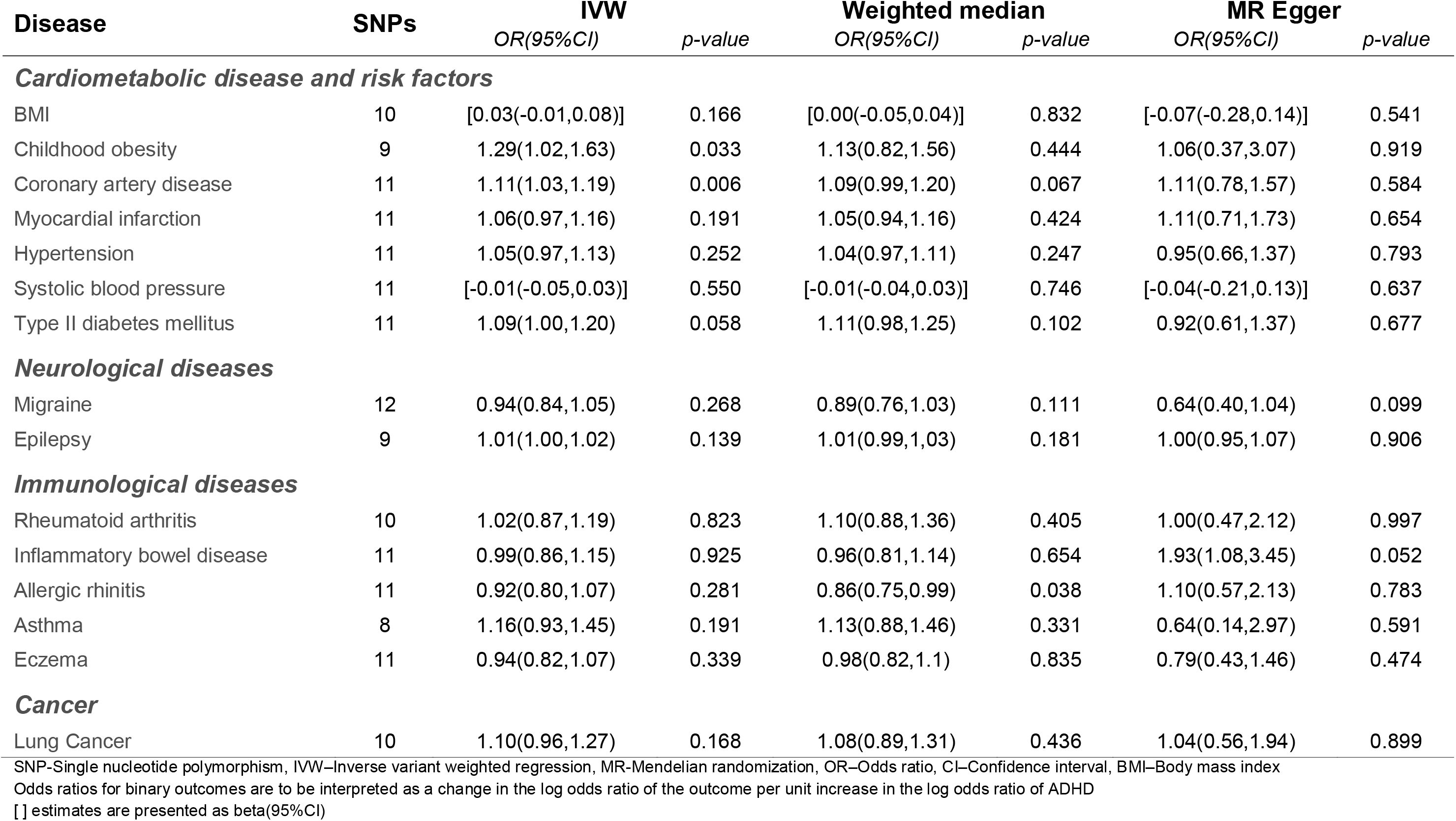
Causal estimates for ADHD on physical diseases using IVW, weighted median and MR Egger regression.

SNPs associated with ADHD were extracted from the respective outcome GWAS (Table S3). If SNPs could not be extracted (due to differences in genotype chips, reference panels or imputation methods between studies), we searched for proxies using SNiPA with r^2^=0.9 (https://snipa.helmholtz-muenchen.de/snipa3/).

### Assessing instrument strength

Instrument strength (assumption (1) of MR) was assessed using the F-statistic, which is equal to the ratio of the mean square of the model estimates to the mean square of the model error for each genetic variant. If F>10, as a rule of thumb, results should not suffer from weak instrument bias(27).

### Estimating causal effects using a two-sample MR approach

The SNP-exposure and SNP-outcome associations were assessed using an inversevariance weighted approach (IVW), a weighted median approach and MR-Egger regression. All three methods require different assumptions on instrument validity to be true and should be interpreted alongside each other. IVW assumes all instruments are valid with no horizontal pleiotropy and therefore restricts the regression intercept to zero (28). Weighted median provides a causal estimate if at least 50% of instruments can be considered valid (29). MR-Egger relaxes the assumption that the outcome is affected only via the exposure and includes a random intercept coefficient in the weighted regression, which reduces power compared to the other two methods. The intercept coefficient is then equivalent to the overall pleiotropic effect and the slope coefficient displays the causal estimate under the assumption that the pleiotropic effects of the genetic variants on the outcome are not proportional to the variants’ effects on the risk factor of interest (30).

Odds ratios (OR) for associations between binary exposures and binary outcomes in two-sample MR studies are interpreted as the OR for outcome per unit increase in the log odds ratio of the exposure.

We present multiple testing uncorrected estimates (while acknowledging the number of correlated phenotypes that have been tested) and focus on consistent results across MR methods to assess the strength of evidence for a causal effect.

### Assessing heterogeneity

Cochrane’s Q test was used to test for heterogeneity in the estimates from the different instruments in IVW. A large test statistic indicates that the estimated causal effects vary across the different instruments. An adaption of the I^2^ statistics often employed in meta-analysis, called I^2^_GX_, was used to test for heterogeneity in MR Egger regression(31). Low I^2^_GX_ values (<90%) suggest heterogeneity and indicate potential violations of the “no measurement error assumption” (NOME) assumption in MR Egger and lead to regression dilution bias. We also performed SIMEX adjusted MR Egger regression where appropriate (R package SIMEX version 1.7) (31).

### Bidirectional MR

When a potentially causal effect of genetic liability for ADHD on a health outcome was detected, we further investigated the direction of effect in a bidirectional MR analysis. In this case, both the effect of the exposure genetic variants on the outcome and the outcome genetic variants to exposure are assessed to inform about the direction of an effect that could be cross-generational. We identified independent genetic variants for the relevant outcomes (childhood obesity (p<1×10^−6^) (16) and CAD (p<5×10^−8^) (17)) from their respective GWAS summary statistics and used the steps described before to perform a two-sample MR.

### Multivariable MR

Two Multivariable MR (MVMR) analysis were conducted to assess the effects of genetic liability for ADHD together with genetic liability for childhood obesity or lifetime smoking heaviness on CAD (32). MVMR is an extension of MR that can be used to estimate the causal effects of multiple exposures on one outcome simultaneously (Figure 2). MVMR is explained in more detail elsewhere (33).

**Figure 2:**
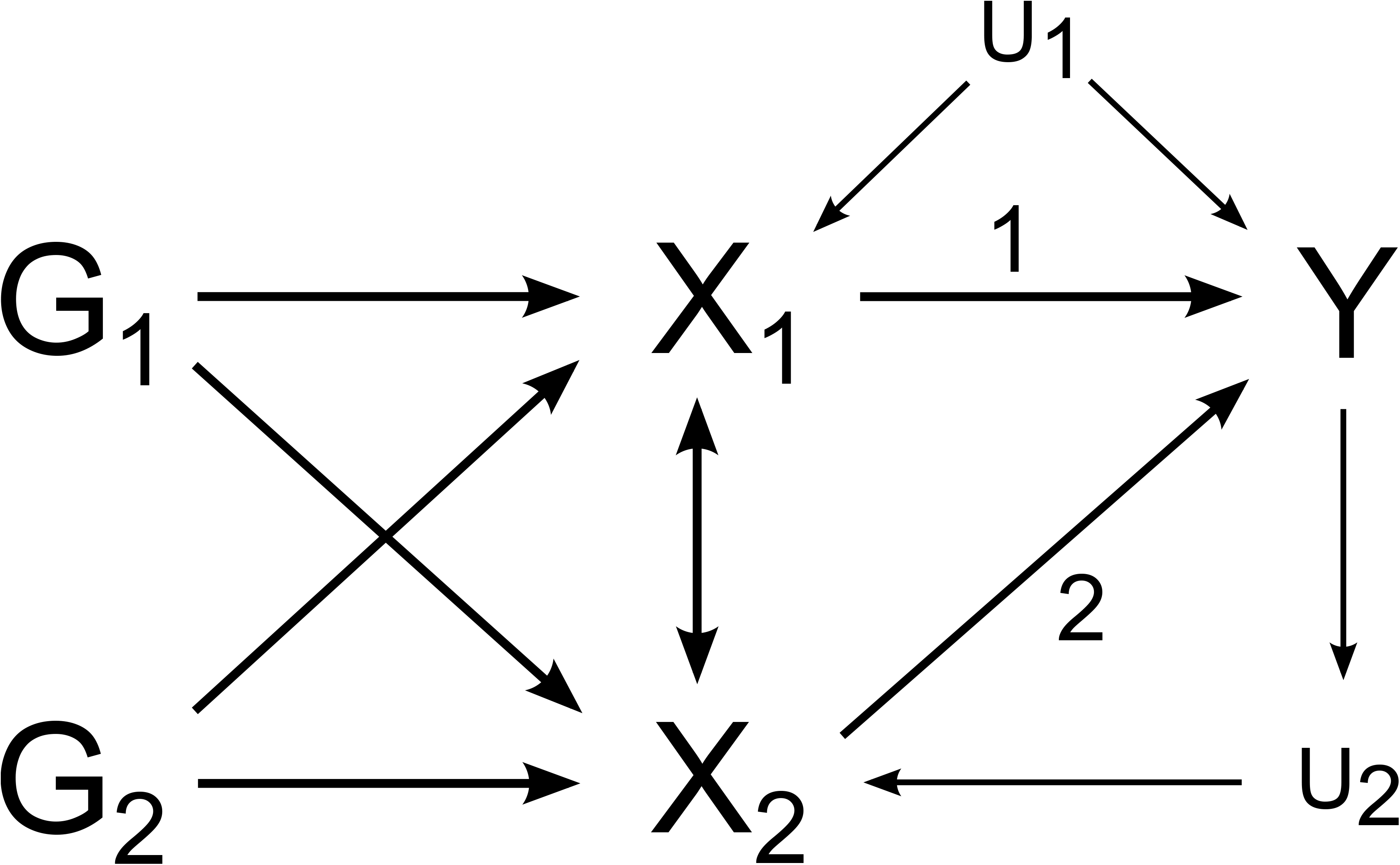
Hypothetical relationship between genetic variants (G), exposures X_1_ and X_2_ and an outcome Y under the presence of unobserved confounders (U).

## RESULTS

### Estimates of causal effects of ADHD on physical health outcomes

#### Cardiometabolic disease and risk factors

Instrument strength for IVW-MR was assessed using F-statistics for the SNP-exposure association (Table S4) and indicated good instrument strength. MR results are shown in Table 1. There was evidence of a causal effect of genetic liability for ADHD on childhood obesity (OR:1.29 (95% CI:1.02,1.63)) and CAD (OR:1.11 (95% CI:1.03,1.19)) using IVW. Both weighted median and MR Egger showed effects consistently in the same direction as IVW although with wider confidence intervals (Figure 3). There was little evidence of horizontal pleiotropic effects (MR Egger intercept for childhood obesity: 1.02 (95% CI:0.92,1.12); for coronary artery disease: 1.00 (95% CI:0.97,1.03)) (Table S5) or heterogeneity between instruments as assessed by Cochrane’s Q statistics (Table S4).

**Figure 3:**
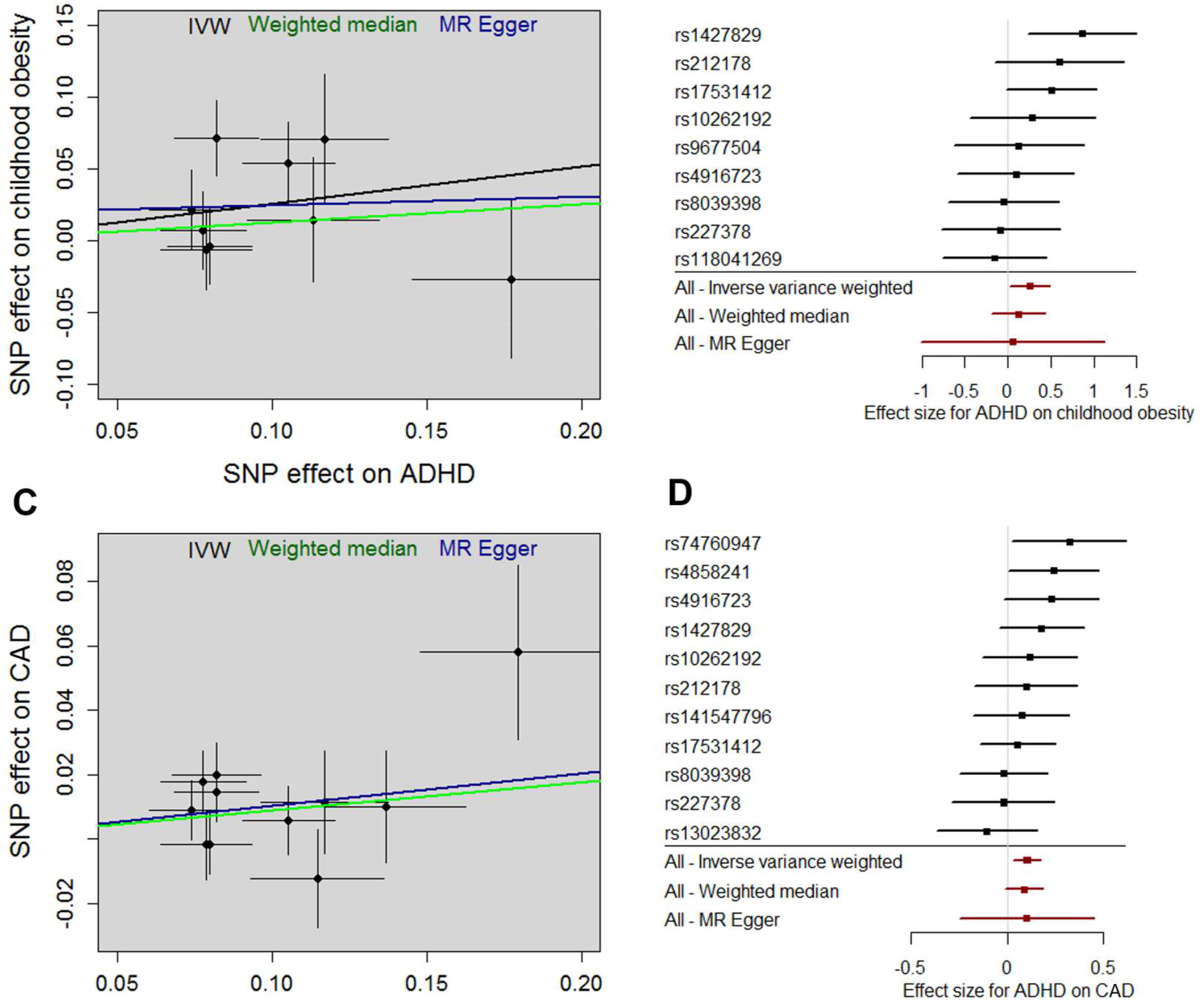
Effect estimates of single variants for ADHD on childhood obesity (A+B) and coronary artery disease (CAD) (C+D) using IVW, weighted median and MR Egger approaches. Estimates are shown as log odds±SE (A+C) and log odds± 95%CI (B+D).

There was little evidence of a causal effect of ADHD genetic liability on other cardiometabolic diseases or risk factors. However, instrument heterogeneity (assessed by Cochrane’s Q statistic) was detected for BMI (p=0.006), hypertension (p=8×10^−5^) and blood pressure (p=8×10^−5^), which might undermine the ability to detect a causal relationship.

#### Neurological diseases

There was little evidence of a causal effect of ADHD genetic liability on migraine (OR:0.94 (95% CI:0.84,1.05)) or epilepsy (OR:1.01 (95% CI:1.00,1.02)).

#### Immunological diseases and cancer

There was little evidence of a causal effect of ADHD genetic liability for immunological diseases and lung cancer from IVW and weighted median estimates. However, horizontal pleiotropy was detected for inflammatory bowel disease (IBD) (MR Egger intercept :0.94 (95% CI:0.89,0.99)) and there was evidence of a causal effect from MR Egger (OR:1.93 (95% CI:1.08,3.45)). There was little evidence of heterogeneity between instruments, except for allergic rhinitis.

Based on these initial findings, there is some evidence of a causal effect of genetic liability for ADHD on childhood obesity, CAD and IBD. Hence, these three disorders were taken forward for further sensitivity analysis.

### Heterogeneity and violation of the NOME assumption

I^2^_GX_ estimates for childhood obesity (I^2^_GX_=47%), CAD (I^2^_GX_=48%) and IBD (I^2^_GX_=48%) (see supplementary Table S4 for all I^2^_GX_ estimates) indicate a potential violation of the NOME assumption which might have induced regression dilution bias in MR-Egger estimates. Hence, we performed MR-SIMEX to adjust MR-Egger estimates accordingly (Table S6). The SIMEX point estimate for CAD was consistent with the original IVW and MR Egger estimate (OR:1.11 (95% CI:1.03,1.20)). For childhood obesity the MR SIMEX estimate (OR:1.26 (95% CI:0.99,1.60)) was increased in comparison to the MR Egger estimate and was closer to the IVW estimate of 1.29. The effect of genetic liability for ADHD on IBD in MR Egger regression was completely attenuated with OR of 1.00 (95% CI:0.86,1.17) after SIMEX adjustment, which is in line with the IVW estimate of 0.99 and hence was not taken forward for further analysis.

### Bidirectional MR for coronary artery disease and childhood obesity

To further investigate the direction of observed effects, we performed bidirectional MR to examine the effect of genetic liability for CAD and childhood obesity on ADHD (Table 2). There was little evidence of a causal effect of genetic liability for CAD on ADHD with an IVW estimate of 0.98 (95% CI:0.92,1.04). However, there was some evidence of a causal association of genetic liability for childhood obesity on ADHD with IVW OR=1.15 (95% CI:1.05,1.25) and weighted median OR=1.10 (95% CI:1.01,1.20). MR Egger estimates were directionally consistent and there was little evidence of heterogeneity or horizontal pleiotropy (MR Egger intercept: 1.00 (95% CI:0.88,1.13)).

**Table 2:** Bidirectional MR with causal estimates for childhood obesity, coronary artery disease and inflammatory bowel disease on ADHD using IVW, weighted median and MR Egger regression.

| Disease | SNPs | IVW |  | Weighted median |  | MR Egger |  |
| --- | --- | --- | --- | --- | --- | --- | --- |
|  |  | OR(95%CI) | p-value | OR(95%CI) | p-value | OR(95%CI) | p-value |
| Childhood obesity | 7 | 1.15(1.05,1.25) | 0.003 | 1.10(1.01,1.20) | 0.031 | 1.16(0.60,2.25) | 0.677 |
| Coronary artery disease | 37 | 0.98(0.92,1.04) | 0.468 | 0.96(0.88,1.05) | 0.346 | 0.91(0.79,1.06) | 0.224 |

SNP - Single nucleotide polymorphism, IVW – Inverse variant weighted regression, MR - Mendelian randomization, OR – Odds ratio, CI – Confidence interval Odds ratios for binary outcomes are to be interpreted as a change in the log odds ratio of the outcome per unit increase in the log odds ratio of the exposure

### Exploratory analyses: Estimates of a causal effect of ADHD on metabolic markers

The two-sample MR of ADHD on metabolic markers suggested little evidence for a causal effect on any of these markers. (Table S7).

### Multivariable MR to test for a potential mediating effect of childhood obesity and lifetime smoking

Obesity and smoking are established risk factors for CAD (34) that are strongly associated with ADHD (35)’(36) and hence may be mediating the association between the genetic liability for ADHD and CAD. When genetic variants for ADHD and childhood obesity were simultaneously entered in the MVMR model, the direct causal effect of the genetic liability for ADHD on CAD was attenuated to 1.06 (95% CI:0.95,1.17), whereas the effect of the genetic liability for childhood obesity on CAD remained stable (OR:1.14(95% CI:1.08,1.20)) (Table S8), which supports a mediating role of childhood obesity. The causal effect of genetic liability for ADHD on lifetime smoking heaviness was 1.07 (95%CI:1.04,1.10). When genetic variants for ADHD and lifetime smoking heaviness were simultaneously entered in the MVMR model, the direct causal effect of the genetic liability for ADHD on CAD remained stable (OR:1.10 (95% CI:1.00,1.21), whereas the effect of the genetic liability for lifetime smoking heaviness on CAD was attenuated (OR:1.38 (95% CI:0.99,1.92)) (Table S9), which supports an effect of genetic liability for ADHD on CAD independent of genetic liability to lifetime smoking heaviness.

## DISCUSSION

In this study, we employed a two-sample MR approach to test for causal effects of genetic liability for ADHD on previously implicated physical health outcomes. By using genetic variants to proxy for ADHD, we found evidence consistent with a causal effect of genetic liability for ADHD on coronary artery disease (CAD) and evidence for a bidirectional association between genetic liability for ADHD and childhood obesity. There was little evidence for a causal effect on neurological and immunological diseases. The causal effect of genetic liability for ADHD on CAD appears to be (at least partially) mediated by childhood obesity and independent of lifetime smoking.

As reported previously ADHD and genetic liability for ADHD are associated with unhealthy lifestyle and risk behaviours in observational studies (12, 37–39). Patients with ADHD are more likely to smoke (35), be overweight (36, 40) and lead a sedentary life (39). All these ADHD outcomes are also risk factors for CAD (34) and hence may act additively as mediators of the observed effect of genetic liability for ADHD on CAD. There are some reports of observational associations between ADHD and cardio-vascular diseases (41, 42), although such studies are difficult to conduct due to the latency between exposure and outcome. Here we report evidence supporting a causal effect of genetic liability for ADHD on CAD but not on other cardiovascular risks (such as hypertension). This effect was attenuated when simultaneously assessing the effect of genetic liability for ADHD and childhood obesity on CAD, suggesting childhood obesity as a potential mediator. However, it remained unchanged when adding genetic liability for lifetime smoking, suggesting an effect of genetic liability for ADHD on CAD independent of genetic liability for lifetime smoking heaviness.

Many studies have shown associations between ADHD and obesity in adolescents and adulthood (36, 40, 43) but the causality of the association has not been established. Hyperactivity is a hallmark of ADHD and it may therefore appear counterintuitive that patients with ADHD have a higher risk for obesity (44). However, observational studies have shown that those with ADHD have been reported to spend more time watching television (45), suffer more school exclusions, show lower levels of physical activity, poorer motor skills and increased dysregulation of eating behaviour (44). Our bidirectional MR between genetic liability for ADHD and genetic liability for childhood obesity supports an explanation of shared familial risks including genetic risk in the form of pleiotropic genetic variants that are causal for both ADHD and childhood obesity; and shared environmental/ foetal risk factors among ADHD and childhood obesity (43).

Our results further suggest little evidence supporting a causal role of genetic liability for ADHD on neurological and immunological diseases. One explanation for this might be that there is no causal effect and associations found in observational studies were better explained by other factors, such as confounding. For example, socio-economic factors can confound the observational associations between ADHD and later health outcomes as both ADHD and health behaviours are strongly associated with lower socio-economic status, educational attainment and lower income. Another explanation is that these null-findings might have arisen due to some of the study limitations, such as instrument validity and population stratification, which are discussed in more detail below.

### Limitations

Since the latest ADHD GWAS was made available in November 2017, there are genetic variants that are strongly associated with ADHD but still only explain little variation in the ADHD phenotype (12). The low I^2^_GX_ estimates suggest substantial heterogeneity in MR Egger regression which is a marker of measurement error in the instruments. This indicates that there might not have been enough power to detect causal associations, since weak instruments bias associations towards the null in two-sample MR studies (27). Furthermore, although there was little statistical evidence for horizontal pleiotropy, the underlying biological pathways leading to ADHD are unknown for most of the genetic variants and therefore the possibility of pleiotropic effects of these variants and hence violation of the exclusion restriction assumption cannot be discounted. Since this analysis employed a two-sample MR framework, which is based on publicly available data, we were not able to test whether the genetic variants used as instruments are independent of potential confounders of the observed exposure to outcome associations. Confounding may also arise due to population stratification (GWAS sample not representative of the underlying population or GWAS samples of mixed populations)(46), assortative mating (traits not inherited independently and consequent violation of the MR assumption that genetic variants are allocated randomly at conception)(47) or selection bias in the GWAS used, all of which might affect both positive and negative findings of our analyses. Furthermore, estimated two-sample MR odds ratios (OR) for associations of binary exposures with binary outcomes can be biased and should only be interpreted in terms of direction and strength of association. (48)

Although MR can overcome some limitations of observational studies, it also makes strong assumptions that cannot always be tested. We therefore highly encourage triangulation of evidence using different causally informative study designs, such as negative control and twin studies (49).

### Conclusion

We found evidence in favour of a causal effect of genetic liability for ADHD on CAD that is potentially mediated by childhood obesity. This strengthens the argument for early treatment and support for children with ADHD and their families and especially promoting physical activity and providing them with dietary advice and support to reduce the future risk for developing obesity and CAD.

## ACKNOWLEGDEMENTS

We are thankful to the PGC, iPSYCH, GIANT, EGG, CARDIoGRAMplusC4D, ILAE, IIBDGC, GABRIEL and ILCCO consortia for making their GWAS summary stats available. Furthermore, we would like to thank Okada et al., Wootton et al., Scott et al. and the Neale Lab to make their GWAS summary stats available.

## CONTRIBUTIONS

Dr Beate Leppert had full access to all the data in the study and take responsibility for the integrity of the data and the accuracy of the data analysis.

*Concept and design:* Leppert, Ajay Thapar, Anita Thapar, Tilling, Davey Smith, Stergiakouli

*Acquisition, analysis, or interpretation of data:* Leppert, Anita Thapar, Tilling, Davey Smith, Stergiakouli

*Drafting of the manuscript:* Leppert

*Critical revision of the manuscript and important intellectual content:* Riglin, Staley, Dardani, Anita Thapar, Ajay Thapar, Tilling, Davey Smith

*Statistical analysis:* Leppert, Dardani, Staley

*Obtained funding:* Thapar, Stergiakouli, Tilling, Davey Smith

*Administrative, technical, or material support:* Dardani, Staley

*Supervision:* Anita Thapar, Tilling, Davey Smith, Stergiakouli

