## Supplementary material for "ADHD genetic liability and physical health outcomes - A two-sample Mendelian randomization study"

Beate Leppert, Lucy Riglin, Christina Dardani, Ajay Thapar, James R Staley, Kate Tilling, George Davey Smith, Anita Thapar, Evie Stergiakouli

**Content**

Supplementary Table S1: ADHD genetic variants

Supplementary Table S2: GWAS summary statistics for outcome measures

Supplementary Table S3: Genetic variants included in each two-sample MR analysis

Supplementary Table S4: Measures of heterogeneity and instrument strength in MR approaches.

Supplementary Table S5: Pleiotropy in MR Egger regression assessed by the MR Egger intercept

Supplementary Table S6: Causal estimates for ADHD on physical diseases using MR Egger and SIMEX adjusted MR Egger regression.

Supplementary Table S7: Causal estimates for ADHD on metabolic markers using IVW, weighted median and MR Egger regression.

Supplementary Table S8: Multivariable MR for ADHD on coronary artery disease with childhood obesity as covariate.

Supplementary Table S9: Multivariable MR for ADHD on coronary artery disease with adult BMI as covariate.

Approximate transformation of association estimates for binary traits in UK Biobank (Neale) GWAS from the risk difference scale to the log odds scale

**Table S1: ADHD genetic variants**

|  | **SNP** | **effect allele** | **other allele** | **beta** | **SE** | **p-value** |
| --- | --- | --- | --- | --- | --- | --- |
| 1 | rs17531412 | A | G | 0.105 | 0.015 | 1.1E-12 |
| 2 | rs1427829 | A | G | 0.082 | 0.014 | 1.3E-09 |
| 3 | rs8039398 | T | C | -0.080 | 0.014 | 3.0E-09 |
| 4 | rs4858241 | T | G | 0.082 | 0.014 | 8.2E-09 |
| 5 | rs28411770 | T | C | 0.086 | 0.015 | 1.2E-08 |
| 6 | rs212178 | A | G | -0.117 | 0.021 | 1.2E-08 |
| 7 | rs74760947 | A | G | -0.180 | 0.032 | 1.4E-08 |
| 8 | rs4916723 | A | C | -0.078 | 0.014 | 1.8E-08 |
| 9 | rs1222063 | A | G | 0.096 | 0.017 | 3.1E-08 |
| 10 | rs10262192 | A | G | 0.074 | 0.014 | 3.7E-08 |
| 11 | rs227378 | A | C | 0.079 | 0.015 | 6.5E-08 |
| 12 | rs13023832 | A | G | 0.115 | 0.022 | 9.3E-08 |
| 13 | rs141547796 | A | G | -0.137 | 0.026 | 9.6E-08 |

**Table S2.** GWAS summary statistics for outcome measures

|  | ***GWAS*** | ***Year*** | ***Ancestry*** | ***Cohort/Consortium*** | ***N_case_*** | ***N_control_*** |
| --- | --- | --- | --- | --- | --- | --- |
| ***Cardiometabolic disease and risk factors*** | | | | | | |
| BMI | Locke et al. | 2015 | European | GIANT | - | 322 154 |
| Childhood obesity | Bradfield et al. | 2012 | European | EGG | 5 530 | 8 318 |
| Coronary artery disease | Nikpay et al. | 2015 | Mixed | CARDIoGRAMplusC4D | 60 801 | 123 504 |
| Myocardial infarction | Nikpay et al. | 2015 | Mixed | CARDIoGRAMplusC4D | 43 676 | 128 199 |
| Hypertension | Neale | 2017 | European | UKBB | 87 690 | 249 469 |
| Systolic blood pressure | Neale | 2017 | European | UKBB | - | 317 754 |
| Type II diabetes mellitus | Scott et al. | 2017 | European | 18 studies^1^ | 26 676 | 132 532 |
| ***Neurological diseases*** | | | | | | |
| Migraine | Neale | 2017 | European | UKBB | 10 007 | 327 152 |
| Epilepsy | ILAE | 2018 | European | ILAE | 15 212 | 29 677 |
| ***Immunological diseases*** | | | | | | |
| Rheumatoid arthritis | Okada et al. | 2014 | European | 18 studies^2^ | 14 361 | 43 923 |
| Inflammatory bowel disease | Liu et al. | 2015 | European | IIBDGC | 12 882 | 21 770 |
| Allergic rhinitis | Neale | 2017 | European | UKBB | 18 934 | 64 595 |
| Asthma | Moffatt et al. | 2010 | European | GABRIEL | 10 365 | 16 110 |
| Eczema | Paternoster et al. | 2015 | European | EAGLE | 10 788 | 30 047 |
| ***Cancer*** | | | | | | |
| Lung Cancer | Wang et al. | 2014 | European | ILCCO | 11 348 | 15 861 |

GWAS – Genome wide association study, BMI- Body mass index

All estimates are assessed in log odds, expect for BMI which was assessed in SD(kg/m2)

1- ARIC, BioMe, deCODE, DGDG, DGI, EGCUT-370, EGCUT-OMNI, EPIC-InterAct, FHS, FUSION, GoDARTS, HPFS, KORAgen, NHS, PIVUS, RS-I, ULSAM, WTCCC

2-BRASS, CANADA, EIRA, NARAC1, NARAC2, WTCCC, Rheumatoid Arthritis Consortium International for Immunochip (RACI)-UK, RACI-US, RACI-SE-E, RACI-SE-U, RACI-NL, RACI-ES, RACI-i2b2, ReAct, Dutch (including AMC, BeSt, LUMC and DREAM), anti-TNF response to therapy collection (ACR-REF: BRAGGSS, BRAGGSS2, ERA, KI and TEAR), CORRONA, Vanderbilt

**Table S3: Genetic variants included in each two-sample MR analysis.**

|  | **independent SNPs associated with ADHD at p<10^-7^** | | | | | | | | | | | | | |
| --- | --- | --- | --- | --- | --- | --- | --- | --- | --- | --- | --- | --- | --- | --- |
|  | rs10262192 | rs1222063 | rs13023832 | rs141547796 | rs1427829 | rs17531412 | rs212178 | rs227378 | rs28411770 | rs4858241 | rs4916723 | rs74760947 | rs8039398 | rs11591402 |
| ***Cardiometabolic disease and risk factors*** | | | | | | | | | | | | | | |
| BMI |  |  | rs9677504 |  |  | rs12410155 | rs12924285 | rs223512 |  |  |  | rs2609653 |  |  |
| Coronary Heart Disease |  |  |  |  |  |  |  |  |  |  |  |  |  |  |
| Childhood obesity |  |  | rs9677504 |  |  | rs12410155 | rs12924285 | rs223512 |  |  |  | rs2609653 |  |  |
| Myocard infarction |  |  |  |  |  |  |  |  |  |  |  |  |  |  |
| Hypertension |  |  | rs9677504 |  |  |  |  | rs150900 |  |  |  |  |  |  |
| Type II Diabetes mellitus |  |  |  |  |  |  |  |  |  |  |  |  |  |  |
| Systolic Blood pressure |  |  | rs9677504 |  |  |  |  | rs150900 |  |  |  |  |  |  |
| ***Psychiatric and Neurological diseases*** | | | | | | | | | | | | | | |
| Migraine |  |  | rs9677504 |  |  |  |  | rs150900 |  |  |  |  |  | rs6584649 |
| Epilepsy |  |  | rs9677504 |  |  |  |  |  |  |  |  |  |  |  |
| Suicidal attempts |  |  | rs9677504 |  |  |  |  | rs150900 |  |  |  |  |  |  |
| ***Inflammatory Diseases*** | | | | | | | | | | | | | | |
| Rheumatoid Arthritis |  |  |  | rs56068671 |  |  |  |  |  |  |  | rs2609653 |  |  |
| Inflammatory Bowel Disease |  |  |  |  |  |  |  |  |  |  |  |  |  |  |
| Allergic rhinitis |  |  | rs9677504 |  |  |  |  | rs150900 |  |  |  |  |  |  |
| Asthma | rs2106900 |  | rs1912185 |  | rs704061 | rs12410155 |  | rs223504 |  |  |  |  | rs281320 |  |
| Eczema |  |  |  |  |  |  |  |  |  |  |  |  |  |  |

Green = SNP extracted, Yellow = proxy SNP extracted, Black = SNP not available, Red = SNP palindromic and excluded

**Table S4: Measures of heterogeneity and instrument strength in MR approaches.**

|  | **F-statistic** | **Cochranes Q** | **I^2^_GX_** |
| --- | --- | --- | --- |
| **Disease** | SNP^2^-ADHD | p-value^1^ |  |
| ***Cardiometabolic disease and risk factors*** | | | |
| BMI [SD(kg/m^2^)] | 33.87 | 0.006 | 0.42 |
| Childhood obesity | 33.96 | 0.318 | 0.47 |
| Coronary artery disease | 33.46 | 0.472 | 0.48 |
| Myocardial infarction | 33.46 | 0.234 | 0.48 |
| Hypertension | 33.45 | 8x10^-5^ | 0.48 |
| Systolic blood pressure | 33.95 | 2x10^-5^ | 0.5 |
| Type II diabetes mellitus | 33.46 | 0.934 | 0.48 |
| ***Psychiatric and Neurological diseases*** | | | |
| Migraine | 33.3 | 0.385 | 0.43 |
| Epilepsy | 34.49 | 0.80 | 0.48 |
| Suicidal behaviour | 33.45 | 0.411 | 0.48 |
| ***Inflammatory Diseases*** | | | |
| Rheumatoid arthritis | 33.72 | 0.362 | 0.5 |
| Inflammatory bowel disease | 33.46 | 0.265 | 0.48 |
| Allergic rhinitis | 33.45 | 0.003 | 0.48 |
| Asthma | 33.61 | 0.151 | 0 |
| Eczema | 33.46 | 0.478 | 0.48 |
| ***Cancers*** | | | |
| Lung Cancer | 33.89 | 0.576 | 0.52 |

1 – for IVW

2 – Variants associated with ADHD that could be extracted from the corresponding disease GWAS summary statistics

**Table S5: Pleiotropy in MR Egger regression assessed by the MR Egger intercept**

| **Disease** | **SNPs** | **MR Egger intercept** | |
| --- | --- | --- | --- |
|  |  | *OR(95%CI)* | *p-value* |
| ***Cardiometabolic disease and risk factors*** | | | |
| BMI [SD(kg/m^2^)] | 10 | 1.01(0.99,1.03) | 0.354 |
| Childhood obesity | 9 | 1.02(0.92,1.12) | 0.717 |
| Coronary artery disease | 11 | 1.00(0.97,1.03) | 0.994 |
| Myocardial infarction | 11 | 1.00(0.96,1.04) | 0.848 |
| Hypertension | 11 | 1.00(1.00,1.01) | 0.604 |
| Systolic blood pressure | 11 | 1.00(0.99,1.02) | 0.722 |
| Type II diabetes mellitus | 11 | 1.02(0.98,1.06) | 0.396 |
| ***Neurological diseases and risk behaviour*** | | | |
| Migraine | 12 | 1.00(1.00,1.00) | 0.140 |
| Epilepsy | 9 | 1.00(1.00,1.01) | 0.848 |
| Suicidal behaviour | 11 | 1.00(1.00,1.00) | 0.366 |
| ***Inflammatory Diseases*** | | | |
| Rheumatoid arthritis | 10 | 1.00(0.93,1.08) | 0.965 |
| Inflammatory bowel disease | 11 | 0.94(0.89,0.99) | 0.046 |
| Allergic rhinitis | 11 | 1.00(0.99,1.01) | 0.606 |
| Asthma | 8 | 1.05(0.92,1.20) | 0.473 |
| Eczema | 11 | 1.02(0.96,1.07) | 0.594 |
| ***Cancer*** | | | |
| Lung Cancer | 10 | 1.01(0.95,1.07) | 0.856 |

**Table S6: Causal estimates for ADHD on physical diseases using MR Egger and SIMEX adjusted MR Egger regression.**

|  |  | **MR Egger** | | | | **SIMEX adjusted MR Egger** | | | |
| --- | --- | --- | --- | --- | --- | --- | --- | --- | --- |
| **Disease** | **SNPs** | **slope** | | **intercept** | | **slope** | | **intercept** | |
|  |  | *OR(95%CI)* | *p-value* | *OR(95%CI)* | *p-value* | *OR(95%CI)* | *p-value* | *OR(95%CI)* | *p-value* |
| Childhood obesity | 9 | 1.06(0.37,3.07) | 0.919 | 1.02(0.92,1.12) | 0.717 | 1.26(0.99,1.60) | 0.102 | 1.01(0.99,1.04) | 0.258 |
| Coronary Heart disease | 11 | 1.11(0.78,1.57) | 0.584 | 1.00(0.97,1.03) | 0.994 | 1.11(1.03,1.20) | 0.028 | 1.00(0.99,1.01) | 0.852 |
| Inflammatory bowel disease | 11 | 1.93(1.08,3.45) | 0.052 | 0.94(0.89,0.99) | 0.046 | 1.00(0.86,1.17) | 0.992 | 1.00(0.98,1.01) | 0.713 |

**Table S7: Causal estimates for ADHD on metabolic markers using IVW, weighted median and MR Egger regression.**

| **Trait** | **Sample Size** | **IV-SNPs** | **IVW** | | | **Weighted median** | | | **MR Egger slope** | | | **MR Egger intercept** | | | **Cochrane’s Q** | **I^2^GX** |
| --- | --- | --- | --- | --- | --- | --- | --- | --- | --- | --- | --- | --- | --- | --- | --- | --- |
|  |  |  | **log odds** | **SE** | **p** | **log odds** | **SE** | **p** | **log odds** | **SE** | **p** | **log odds** | **SE** | **p** |  |  |
| Omega-3 fatty acids | 13544 | 11 | 0.077 | 0.046 | 0.095 | 0.093 | 0.060 | 0.120 | 0.276 | 0.182 | 0.163 | -0.020 | 0.017 | 0.286 | 0.646 | 0.484 |
| Omega-6 fatty acid | 13506 | 11 | 0.052 | 0.046 | 0.255 | 0.040 | 0.058 | 0.488 | -0.039 | 0.182 | 0.835 | 0.009 | 0.017 | 0.617 | 0.922 | 0.484 |
| Total cholesterol | 21491 | 11 | 0.043 | 0.038 | 0.251 | 0.071 | 0.049 | 0.150 | -0.075 | 0.153 | 0.638 | 0.011 | 0.014 | 0.448 | 0.908 | 0.484 |
| Serum total triglycerids | 21545 | 11 | 0.055 | 0.040 | 0.171 | 0.058 | 0.053 | 0.268 | 0.044 | 0.174 | 0.805 | 0.001 | 0.016 | 0.950 | 0.307 | 0.484 |
| Total fatty acids | 13505 | 11 | 0.066 | 0.046 | 0.152 | 0.044 | 0.058 | 0.448 | 0.072 | 0.182 | 0.704 | -0.001 | 0.017 | 0.975 | 0.956 | 0.484 |
| Total cholesterol in LDL | 21559 | 11 | 0.049 | 0.037 | 0.190 | 0.069 | 0.047 | 0.137 | -0.097 | 0.152 | 0.541 | 0.014 | 0.014 | 0.350 | 0.890 | 0.484 |
| Total cholesterol in HDL | 21555 | 11 | -0.023 | 0.037 | 0.544 | -0.039 | 0.050 | 0.434 | -0.199 | 0.153 | 0.225 | 0.017 | 0.014 | 0.265 | 0.588 | 0.484 |
| Free cholesterol | 13497 | 11 | 0.009 | 0.049 | 0.849 | 0.037 | 0.066 | 0.577 | 0.013 | 0.204 | 0.951 | 0.000 | 0.019 | 0.986 | 0.334 | 0.484 |

**Table S8: Multivariable MR for ADHD on coronary artery disease with genetic liability for childhood obesity as covariate.**

|  |  | **IVW** | | | | | | |
| --- | --- | --- | --- | --- | --- | --- | --- | --- |
|  |  | **Single IV** | | |  | **Multivariable** | | |
|  | **SNP** | **OR** | **95% CI** | **p-value** | **SNP** | **OR** | **95% CI** | **p-value** |
| ADHD | 9 | 1.10 | 1.01,1.19 | 0.022 | 16 | 1.06 | 0.95,1.17 | 0.310 |
| Childhood obesity | 7 | 1.15 | 1.08,1.23 | 9.2x10-6 |  | 1.14 | 1.08,1.20 | 2.5x10-4 |

**Table S9: Multivariable MR for ADHD on coronary artery disease with genetic liability for lifetime smoking as covariate.**

|  |  | **IVW** | | | | | | |
| --- | --- | --- | --- | --- | --- | --- | --- | --- |
|  |  | **Single IV** | | |  | **Multivariable** | | |
|  | **SNP** | **OR** | **95% CI** | **p-value** | **SNP** | **OR** | **95% CI** | **p-value** |
| ADHD | 9 | 1.19 | 1.09,1.29 | 6.1x10-5 | 134 | 1.10 | 1.00,1.21 | 0.045 |
| Lifetime smoking | 130 | 1.69 | 1.30,2.19 | 8.6x10-5 |  | 1.38 | 0.99,1.92 | 0.061 |

Approximate transformation of association estimates for binary traits in UK Biobank (Neale) GWAS from the risk difference scale to the log odds scale :

log_OR = beta/mu(1-mu)

se_log_OR = se/mu(1-mu)

where mu=n_cases/n_total.
